# Alpha-synuclein knockout impairs melanoma development and alters DNA damage repair in the TG3 mouse model in a sex-dependent manner

**DOI:** 10.1101/2024.12.01.626256

**Authors:** Moriah R. Arnold, Suzie Chen, Vivek K. Unni

## Abstract

Strong evidence suggests links between Parkinson’s Disease (PD) and melanoma, as studies have found that people with PD are at an increased risk of developing melanoma and those with melanoma are at increased risk of developing PD. Although these clinical associations are well-established, the cellular and molecular pathways linking these diseases are poorly understood. Recent studies have found a previously unrecognized role for the neurodegeneration-associated protein alpha-synuclein (αSyn) in melanoma; the overexpression of αSyn promotes melanoma cell proliferation and metastasis. However, to our knowledge, no studies have investigated the role of αSyn in *in vivo* melanoma models outside of a xenograft paradigm. Our study created and characterized *Snca* knockout in the spontaneously developing melanoma TG3 mouse line, TG3+/+*Snca*-/-. We show that αSyn loss-of-function significantly delays melanoma onset and slows tumor growth *in vivo*. Furthermore, decreased tumor volume is correlated with a decreased DNA damage signature and increased apoptotic markers, indicating a role for αSyn in modulating the DNA damage response (DDR) pathway. Overall, our study provides evidence that targeting αSyn and its role in modulating the DDR and melanomagenesis could serve as a promising new therapeutic target.

## INTRODUCTION

The association between Parkinson’s Disease (PD) and melanoma has been well established. Many epidemiological studies have found a significant increase in the risk of melanoma among individuals with PD compared to healthy individuals, ranging from 1.4-20- fold (Bajaj et al., 2010, Becker et al., 2010, Bertoni et al., 2010, Constantinescu et al., 2014, Dalvin et al., 2017, Driver et al., 2007, Elbaz et al., 2002b, Kareus et al., 2012, Lo et al., 2010, Moller et al., 1995, Olsen et al., 2006, Olsen et al., 2005, Olsen et al., 2007, Rugbjerg et al., 2012, Ryu et al., 2020, Schwid et al., 2010, Zhang et al., 2021). Likewise, there is also an increased risk for PD in melanoma patients, ranging from 1.7-4.2 fold (Baade et al., 2007, Dalvin et al., 2017, Freedman et al., 2005, Gao et al., 2009, Kareus et al., 2012). Altogether, it is clear that common environmental, genetic, and/or molecular mechanisms are at play to influence this clinical association, yet the underlying mechanism is still poorly understood.

One potentially promising avenue of investigation is the biological function of the neurodegeneration-associated protein, alpha-synuclein (αSyn). Misfolded and aggregated forms of αSyn are found in cytoplasmic inclusions called Lewy bodies, which are neuropathological hallmarks in PD and other Lewy body disorders (Baba et al., 1998, Spillantini et al., 1998). Lewy bodies are found primarily in the central nervous system, where their presence in dopaminergic neurons in the midbrain is associated with the degeneration of these cells in PD (Xu et al., 2002). αSyn is not only found in the central nervous system, but can also be found in the periphery, including in melanocytes (Fayyad et al., 2019, Ma et al., 2019) and therefore could be a key molecular link between these disease pathologies. In primary and metastatic melanoma, ∼85% of biopsies show high expression of αSyn (Arnold et al., 2024, Matsuo and Kamitani, 2010, Rodriguez-Leyva et al., 2017, Rugbjerg et al., 2012, Welinder et al., 2014). Since this initial characterization, there have been several studies investigating the role of αSyn in melanoma growth and metastasis; the majority of these being *in vitro* studies. Overall, these studies using human and mouse melanoma cell lines have found that αSyn expression is important in cell proliferation (Arnold et al., 2024, Israeli et al., 2011, Shekoohi et al., 2021, Turriani et al., 2017), motility (Gajendran et al., 2023), and protects against cell death (Nandakumar et al., 2017, Turriani et al., 2017), through multiple potential mechanisms, such as altering the inflammatory response (Aloy et al., 2024, Fokken et al., 2024, Rajasekaran et al., 2024), autophagy pathways (Nandakumar et al., 2017, Turriani et al., 2017), and DNA damage repair (Arnold et al., 2024).

Fewer studies have investigated the role of αSyn in *in vivo* melanoma mouse models and all this previous *in vivo* work, to our knowledge, has used a xenograft paradigm. In general, these xenograft studies corroborate previous *in vitro* work and find that αSyn is important in melanoma tumor growth and metastasis. Specifically, αSyn knockout (KO) human/mouse melanoma cells implanted as xenografts in mice exhibited slower growth and increased apoptosis (Shekoohi et al., 2021), and reduced tumor-induced mechanical allodynia (Niederberger et al., 2023). Furthermore, WT melanoma cells in αSyn overexpressing mice show increased metastasis (Israeli et al., 2011). Lastly, human melanoma xenografts implanted in mice and treated with an αSyn aggregation inhibitor (anle138b) led to increased cell death (Turriani et al., 2017) and upregulation of anti-melanoma immune responses (Fokken et al., 2024). Despite this substantial data linking αSyn to melanoma tumor growth *in vivo*, whether αSyn expression within melanocytes influences tumorigenesis is still not understood. In our current study, we aimed to create and characterize a new TG3 *Snca*-/- mouse line to better understand the function of αSyn in melanomagenesis, tumor growth, and metastasis in a spontaneous melanoma-forming mouse line. TG3 mice display melanin-pigmented lesions after a short latency and with complete penetrance (Chen et al., 1996, Pollock et al., 2003, Zhu et al., 2000, Zhu et al., 1998). This model is driven by multiple tandem insertions of a transgene into intron 3 of *Grm1* (metabotropic glutamate receptor 1) with concomitant deletion of an intronic sequence that increases expression of *Grm1*. Homozygous TG3 mice form primary melanoma tumors on pinna and perianal regions, in addition to metastatic tumors in lymph nodes, lung, and liver (Chen et al., 1996, Pollock et al., 2003, Zhu et al., 2000, Zhu et al., 1998).

Our previous work has shown αSyn’s role in modulating nuclear DNA damage response (DDR) pathways in human melanoma cells (Arnold et al., 2024) and other cell types (Rose et al., 2024, Schaser et al., 2019). Specifically, we found a novel function of αSyn in DNA double-strand break (DSB) repair, where αSyn colocalizes with DSB repair components and its knockout leads to increased DSBs and their slowed repair (Arnold et al., 2024, Schaser et al., 2019). In this study, we aimed to investigate whether similar mechanisms are important for melanomagenesis and growth using the TG3+/+*Snca* -/- mouse model to test whether αSyn loss-of-function dysregulated DNA damage pathways and led to downstream cell death phenotypes.

## RESULTS

### Loss of alpha-synuclein delays melanoma onset and decreases tumor growth *in vivo*

To study the role of αSyn in melanoma tumorigenesis *in vivo*, TG3 mice (Pollock et al., 2003) were crossed with *Snca*-knockout mice. The generated TG3+/+*Snca*+/+ (“wildtype”) and TG3+/+*Snca*-/- (“homozygous KO”) mice were then analyzed for tumor growth from P30 to P100, at which point the mice were sacrificed and dissected for tissue processing (Figure 1A). There was no significant difference in weight of the mice between wildtype and homozygous KO genotypes (Figure S1). Melanoma tumor onset was evaluated, and homozygous KO mice developed melanoma significantly later compared to the wildtype control group (Figure 1B). Wildtype mice on average exhibited tumors at P43, whereas melanoma onset was observed on average at P50 for homozygous KO mice. This difference is driven primarily by male mice, since when stratified by sex there was no significant difference between genotypes in female mice but there was in male mice (Figure 1B). Further, the progression of melanoma growth on pinna and perianal regions were followed for ∼70 days. Here, a graded scoring system from minimal (0) to extreme tumor growth (5) was used to quantify melanoma progression on pinna as previously described (Schiffner et al., 2012) (Figure S2) and quantitative size measurement was used to quantify melanoma progression in perianal regions. This analysis revealed no significant differences in tumor progression of the pinna between wildtype and homozygous KO genotypes, even when stratified by sex (Figure 1B). However, homozygous KO mice display decreased perianal tumor growth compared to wildtype mice, which becomes significant at later time points (Figure 1C). Again, this difference is driven primarily by male mice when stratified by sex (Figure 1C).

**Figure 1.**
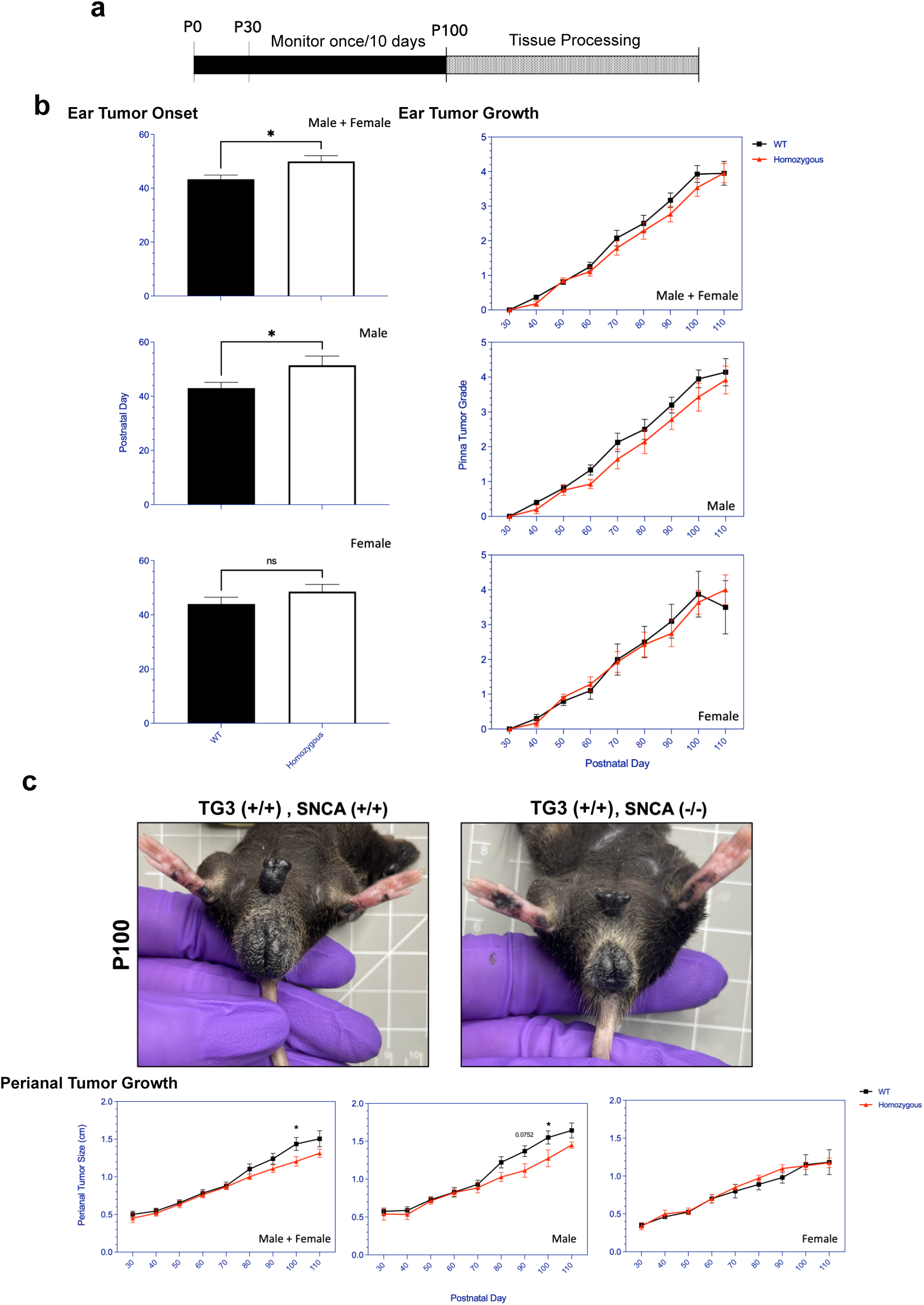
Alpha-synuclein knockout significantly delays tumor onset and slows tumor progression. a) Schematic representing experimental timeline. b,c) Pinna melanoma onset in TG3+/+*Snca*+/+ (n=15) and TG3+/+*Snca*-/- (n=14). Pinna and perianal tumor progression of the TG3+/+*Snca*-/- compared to the TG3+/+*Snca*+/+ after tumor onset. The grading system to evaluate the progression of tumor growth at the pinna region until endpoint at P110 is further described in Figure S2. Analysis was further stratified by sex with TG3+/+*Snca*+/+ male (n=10), TG3+/+*Snca*+/+ female (n=5), TG3+/+*Snca*-/- male (n=7), and TG3+/+*Snca*-/- female (n=7). Error bars represent Standard Error of the Mean (SEM). *p<0.05 by unpaired T-test for tumor onset or Two-way ANOVA for tumor progression.

### Experimental genotypes display similar pigment formation and *Grm1* expression

We next wanted to confirm the presence of melanoma-like cells in the primary tumors of these mice through histopathological analysis. Hematoxylin and eosin (H+E) staining revealed significant levels of pigmented cell growth in the primary pinna and perianal tumors of both wildtype and homozygous KO mice compared to control C57BL/6 wildtype mice without tumors (Figure 2A). qRT-PCR analyses revealed comparable *Grm1* mRNA expression levels between the wildtype and homozygous KO pinna and perianal primary tumors. This suggests that mice express the *Grm1* transgene at similar levels regardless of *Snca* expression (Figure 2B). These levels were compared to positive control cerebellum tissue where *Grm1* mRNA expression is known to be high.

**Figure 2.**
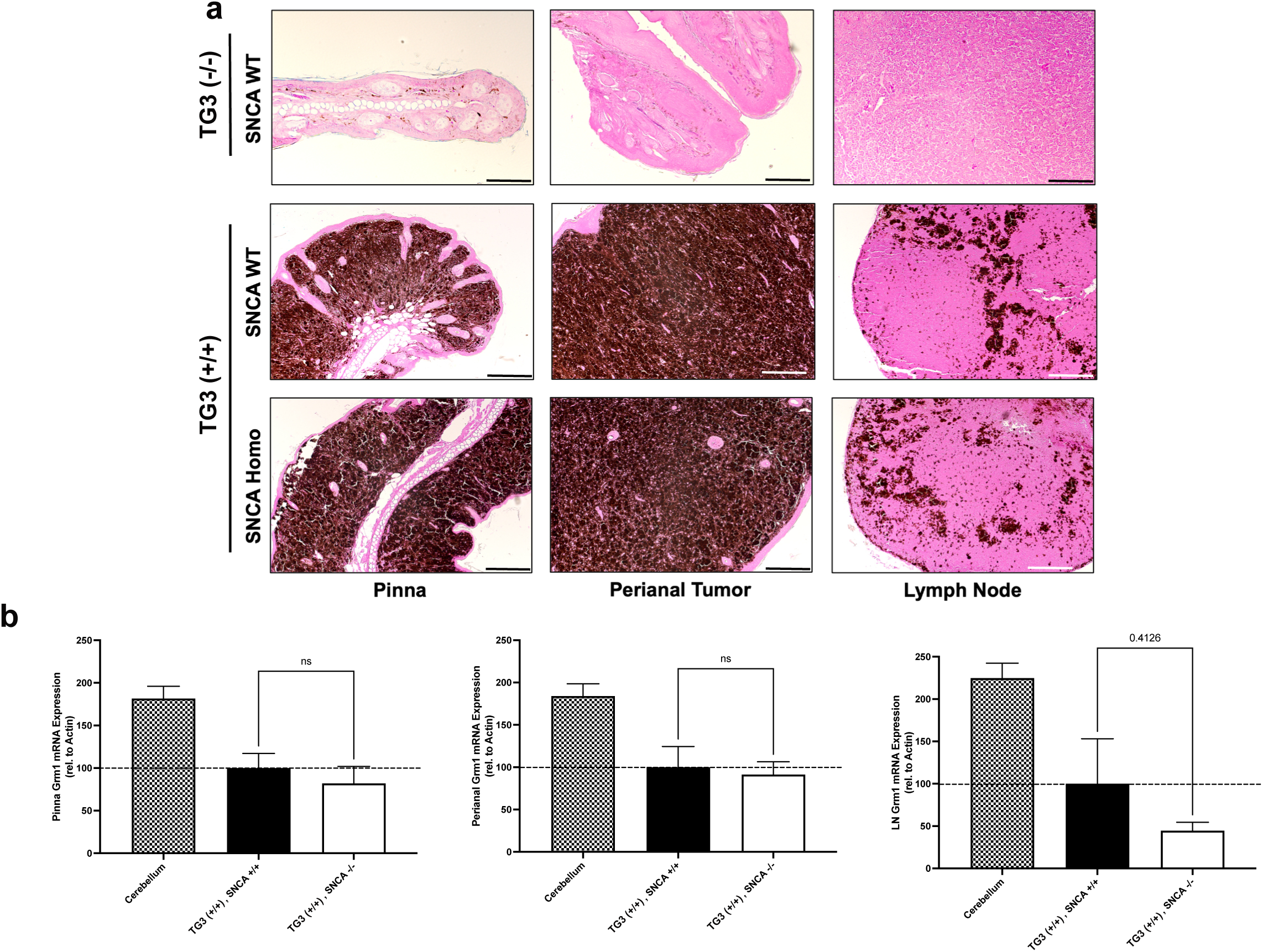
Alpha-synuclein knockout does not interfere with *Grm1* expression and results in possible decrease in lymph node metastasis. a) Formalin-fixed paraffin-embedded pinna and perianal primary tumors and lymph nodes were stained for hematoxylin and eosin in TG3-/-*Snca*+/+, TG3+/+*Snca*+/+, or TG3+/+*Snca*-/- mice. Stained samples were imaged on the Zeiss ApoTome2 microscope. Scale bar=100µm, except for TG3-/-*Snca*+/+ perianal scale bar=200µm and lymph node scale bar=50µm. b) Total RNA was isolated from pinna and perianal primary tumors and lymph nodes from TG3+/+*Snca*+/+ or TG3+/+*Snca*-/- mice. Using primers against the *Grm1* gene, qRT-PCR amplification was determined when normalized to *beta-actin*. For pinna and perianal analysis, TG3+/+*Snca*+/+ (n=5) and TG3+/+*Snca*-/- (n=6). For lymph node analysis, TG3+/+*Snca*+/+ (n=12) and TG3+/+*Snca*-/- (n=9). Cerebellum samples (n=3). Each sample was run with 2 technical replicates. Statistical analysis via unpaired T-test.

Additionally, H+E staining confirmed the presence of pigmented melanoma cells in the lymph nodes of both wildtype and homozygous KO mice, indicating lymph node metastasis had occurred in both genotypes (Figure 2A). The *Grm1* mRNA expression in lymph node tissues of wildtype and homozygous KO mice was analyzed as a marker for melanoma cell dissemination. Here a less, although not significant, *Grm1* expression was observed in lymph nodes of the homozygous KO mice compared to wildtype mice (Figure 2B), suggesting that αSyn loss-of function may decrease metastatic potential but that larger cohorts would be needed to detect this possible difference given inter-animal variability.

### Loss of alpha-synuclein decreases DNA damage signatures in the melanoma tumor

αSyn has been previously linked to DNA DSB repair, since knocking out αSyn significantly increases DNA damage levels in SK-Mel28 cells (Arnold et al., 2024), Hap1 cells (Rose et al., 2024, Schaser et al., 2019), and mouse brain (Schaser et al., 2019) due to less efficient DNA DSB repair. We wanted to investigate whether there were differences in DNA damage burden and repair mechanisms between TG3+/+*Snca* +/+ (“wildtype”) and *Snca* -/- (“homozygous KO”) mice. Formalin-fixed paraffin-embedded perianal primary tumor samples from wildtype and homozygous KO mice underwent immunofluorescence (IF) staining. Genotypes were first validated via IF when stained using an αSyn antibody, LB509, where homozygous KO tissue showed significantly reduced levels of staining compared to wildtype mice (Figure 3A). In addition, when analyzing the localization of αSyn labelling in the wildtype samples, discrete nuclear foci were seen in the melanoma tumor cells, similar to our previous studies where these foci are implicated in DNA damage repair processes (Arnold et al., 2024, Rose et al., 2024, Schaser et al., 2019). Given these findings and previous data, these samples were next stained for DNA damage and damage repair markers: γH2AX, RPA32, and 53BP1.

**Figure 3.**
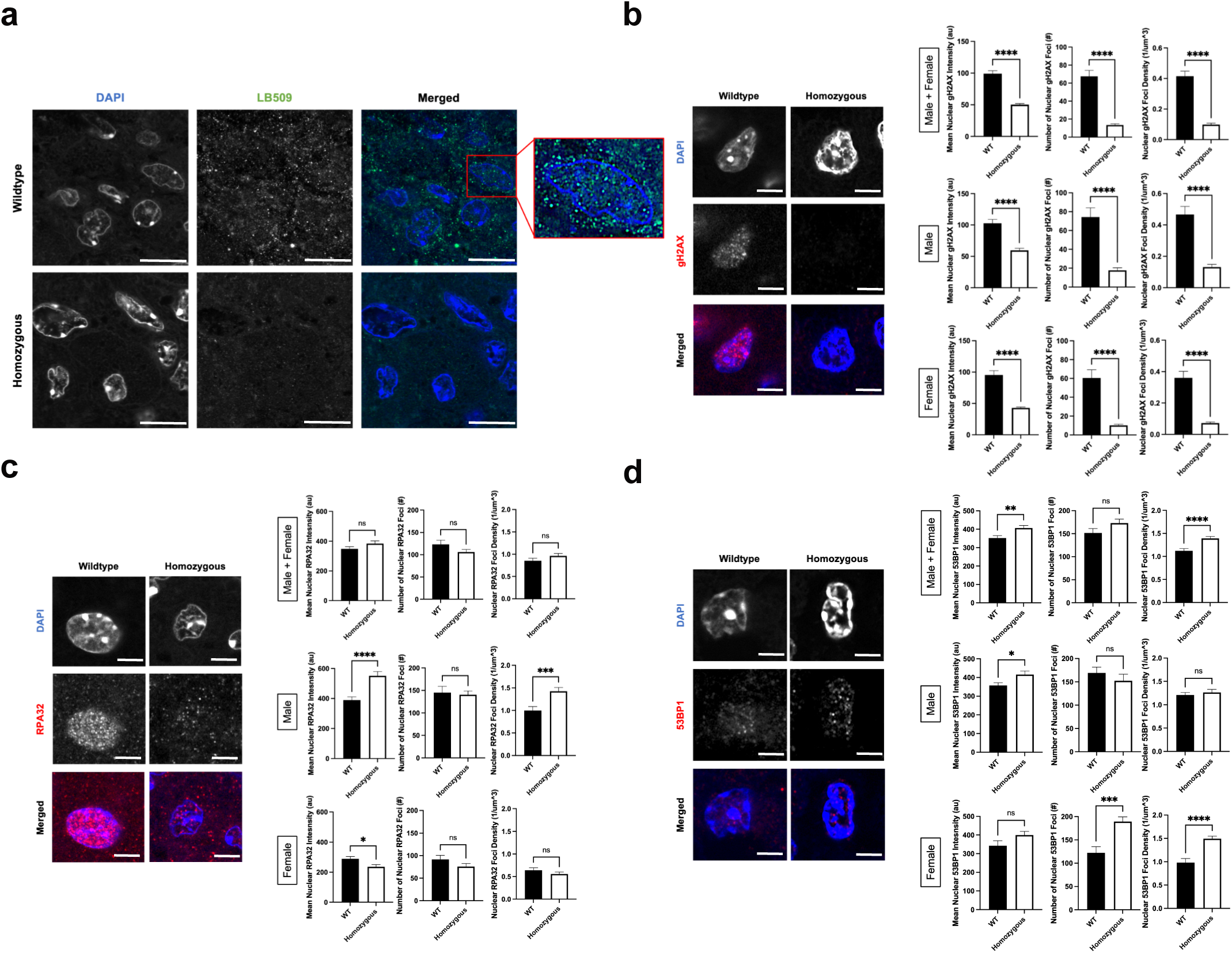
Alpha-synuclein loss-of-function leads to lower DNA damage signature in P110 perianal tumors. a, b, c, d) Formalin-fixed paraffin-embedded perianal primary tumors from TG3+/+*Snca*+/+ (n=5) and TG3+/+*Snca*-/- (n=5) were stained for LB509, _γ_H2AX, RPA32, 53BP1, or DAPI. Stained samples were imaged on the Zeiss 990 confocal microscope with Airyscan processing. Mean intensity, number of foci, and density of foci within DAPI masks were analyzed using Arivis. * p<0.05, ** p<0.01, **** p<0.0001 by unpaired T-test. Error bars denote SEM. Scale bar=5µm (a) or 2µm (b,c,d). Quantification from 5 biological replicates (separate animals) per group were performed (n=163-249 nuclei analyzed per condition).

γH2AX, a phosphorylated form of histone H2AX, is involved in the early stages of DNA DSB detection and is a sensitive marker for DNA damage burden. IF staining for γH2AX revealed a significant decrease in mean intensity of γH2AX signal, number of nuclear γH2AX foci, and density of nuclear γH2AX foci in the homozygous KO group compared to the wildtype group (Figure 3B). These trends remained similar when stratified by sex. RPA32, replication protein A2, binds and stabilizes single-stranded DNA intermediates that form during DNA repair and is important in homologous recombination (HR) DSB repair. IF staining for RPA32 revealed no significant differences in the mean nuclear intensity, number of nuclear RPA32 foci and their density in the homozygous KO group compared to the wildtype group (Figure 3C). Interestingly when stratified by sex, there were significant, but opposite, differences in mean nuclear intensity of RPA32 between the wildtype and homozygous KO group, despite no significant differences when combined. Male homozygous KO mice exhibited a significant increase in mean nuclear RPA32 intensity compared to wildtype mice, whereas female homozygous KO mice exhibited a significant decrease in mean nuclear RPA32 intensity compared to wildtype mice (Figure 3C). Lastly, 53BP1, p53-binding protein 1, is an important regulator of DNA DSB repair and promotes non-homologous end-joining (NHEJ) DSB repair. IF staining for 53BP1 revealed a significant increase in mean nuclear 53BP1 intensity in homozygous KO mice compared to wildtype mice, driven by both male and female mice (Figure 3D). Homozygous KO female mice showed a significant increase in number and density of 53BP1 foci compared to wildtype mice, but male mice showed no genotype differences.

### DNA damage signature correlates to cell death phenotypes

To further understand the downstream cellular consequences of altered DNA damage repair mechanisms in αSyn homozygous KO mice, we assayed various cell death markers via qRT-PCR. These included markers for apoptosis (*Caspase-3*, *Caspase-9*), necroptosis (*RIP3*), autophagic cell death (*LC3B*), and senescence (*Cdkn2a-p16*). We found that perianal tumors of homozygous KO mice exhibited significantly higher gene expression levels of *Caspase-9*, *LC3B*, and *p16* compared to wildtype tumors (Figure 4A). Furthermore, to directly correlate the DNA damage signatures seen with immunofluorescence (Figure 3) with *Caspase-9*, *LC3B*, and *p16*, we ran linear regression analyses. Average nuclear mean intensity, foci number, and foci density of γH2AX and 53BP1 showed no significant associations with these cell death markers (data not shown). However, RPA32 nuclear mean intensity and foci density significantly correlated with the levels of *p16*, with mean foci number close to significance, but did not correlate with *Caspase-9* or *LC3B* levels (Figure 4B). This indicates that mice with higher RPA32 levels also showed higher *p16* mRNA levels, and this trend was statistically significant. This data suggests that αSyn loss-of-function and the subsequent dysregulation of the DDR this causes leads to a senescence-like phenotype, potentially driving the impaired tumor growth we measured *in vivo*.

**Figure 4.**
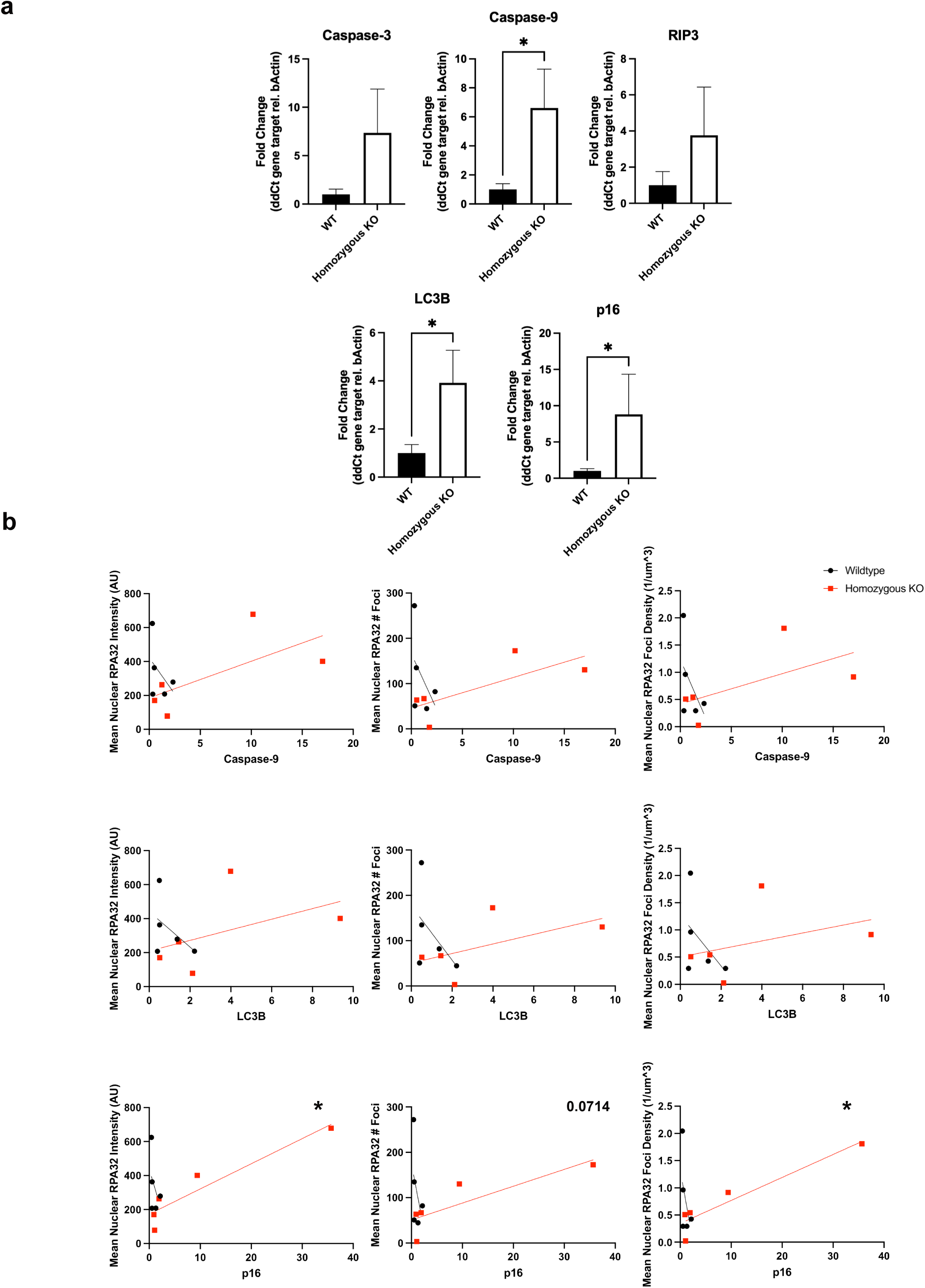
Alpha-synuclein knockout increases apoptosis, autophagy, and senescence marker expression. a) Total RNA was isolated from perianal primary tumors from TG3+/+*Snca*+/+ or TG3+/+*Snca*-/- mice. Using primers against the various genes described in Table S1, qRT-PCR amplification was determined when normalized to *beta-actin* internal control. For analysis, TG3+/+*Snca*+/+ (n=5) and TG3+/+*Snca*-/- (n=6). Each sample was run with 2 technical replicates. * p<0.05 by unpaired Mann-Whitney T-test. b) Simple linear regression analysis of mean nuclear intensity, number of foci, and density of foci of RPA32 immunofluorescence (Figure 3) compared to gene expression of *Caspase-9, LC3B,* and *p16*. Each point represents a single animal, with TG3+/+*Snca*+/+ (n=5, same animals from Figure 3) and TG3+/+*Snca*-/- (n=5, same animals from Figure 3). * p<0.05 by simple linear regression with 95% confidence intervals.

## DISCUSSION

In this study, we extend our knowledge of the molecular connection between PD and melanoma, by uncovering roles for the neurodegeneration-associated protein, αSyn, in melanoma formation and growth. We developed a model to investigate αSyn deficiency on melanomagenesis and metastasis *in vivo* in TG3 mice (Chen et al., 1996, Pollock et al., 2003, Zhu et al., 2000, Zhu et al., 1998). Our data suggest that αSyn loss-of-function significantly delayed melanoma tumor onset in primary pinna tumors and growth of primary perianal tumors. Furthermore, there was a non-significant, but trending, decrease in the metastasis to lymph nodes as measured by *Grm1* mRNA expression. Immunofluorescence staining of the primary perianal tumors revealed a significantly decreased DNA damage signature in *Snca* KO mice, as measured by quantifying nuclear γH2AX. Interestingly, there were sex-dependent differences in nuclear 53BP1 and RPA32 levels in homozygous KO mice compared to wildtype. Lastly, cell death marker analysis revealed that homozygous KO perianal tumors exhibited significantly higher levels of the apoptosis marker *Caspase-9*, autophagic marker *LC3B*, and senescence marker *p16*. In homozygous KO tumors, RPA32 immunofluorescence significantly correlated with *p16* mRNA levels, suggesting a potential senescence-like phenotype partly controlled by dysregulated RPA32-dependent DDR.

These results fit into a larger landscape of links between cancer growth, genomic instability, and DDR. Due to their highly proliferative nature, cancers are especially vulnerable to replication-induced DNA damage and genome instability. Inherently, this leads to the high DNA damage signatures seen in many cancer types (Groelly et al., 2023) and melanoma cells upregulate DSB repair pathway proteins (Puts et al., 2020, Song et al., 2013) to increase metastatic potential (Kauffmann et al., 2008). Our findings suggest that when αSyn is present (“wildtype” mice), DSB repair pathways remain intact, allowing for cell survival and tumor growth. However, DNA damage from hyperproliferation creates large DNA damage signatures in late-stage tumors (Graphical Abstract). In contrast, when αSyn is not present (“homozygous KO” mice), there is impaired DSB repair due to a decrease in DSB repair machinery (Arnold et al., 2024). Accumulation of unrepaired DSBs ultimately leads to cell death and senescence phenotypes, with data suggesting that RPA32 levels synergize with senescence marker *p16* upregulation. This could result in the impaired tumor growth and a decreased DNA damage signature (γH2AX) we detect, since the cells with a high DNA damage signature die and are removed from late-stage tumors (Graphical Abstract). As a consequence of this unrepaired DNA damage and subsequent cell death, remaining cells may upregulate DDR pathways components 53BP1 and RPA32 and this may be sex dependent. Overall, our findings suggest that αSyn upregulation in melanoma may be part of a mechanism to improve DSB repair, allowing cells to evade the programmed cell death that would normally be triggered by high DSB levels, similar to what is seen with the upregulation of other DSB repair pathway proteins (Puts et al., 2020, Song et al., 2013),.

Loss of αSyn resulted in upregulation of various cell death and senescence markers, likely downstream of dysregulated DDR and resulting in the decreased tumor growth *in vivo*. Caspase-9 is an initiator caspase in the intrinsic apoptosis pathway that is activated when cytochrome c is released from mitochondria in response to death signals (Sever et al., 2023). LC3B, microtubule-associated protein 1 light chain-3B, is an autophagic protein that plays a role in cell death and autophagy. Autophagic cell death, also known as type 2 cell death, is characterized by large-scale autophagic vacuolization of the cytoplasm (Kroemer and Levine, 2008, Liu et al., 2023). In general, autophagy can protect cells from stresses like nutrient depletion or starvation, but excessive autophagy can lead to cell death. Furthermore, LC3B can also promote apoptosis through interactions with the extrinsic apoptotic factor Fas (Chen et al., 2010). Lastly, p16(INK4a) is a cyclin-dependent kinase inhibitor that is often expressed in senescent cells, which have stopped growing due to stress (Coppe et al., 2011). This tumor suppressor gene is commonly mutated in human tumors, allowing precancerous lesions to bypass senescence (LaPak and Burd, 2014). These processes have all been associated with DNA damage accumulation and implicated in melanoma, where suppression of Caspase-9 and p16 and over-stimulation of LCB3 have been linked to melanomagenesis, contributing to disease progression and resistance to therapy (Berthenet et al., 2020, Constantinou and Bennett, 2024, Fratta et al., 2023, Gray-Schopfer et al., 2006, Haferkamp et al., 2008, Hartman, 2020, Soengas et al., 2001, Soengas and Lowe, 2003, Szmurlo et al., 2024, Tang et al., 2016). Targeted therapy of some of these modulators is currently being explored as potential therapeutic strategies for melanoma (Fratta et al., 2023, Jaune et al., 2021). Interestingly, our data suggesting a significant synergistic effect of RPA32 protein levels and *p16* expression coincides with previous reports of “RPA exhaustion”. This is a phenomenon by which persistent DNA damage can lead to replication catastrophe and cells then acquire senescent traits and is associated with various age-related pathologies(Feringa et al., 2018, Ibler et al., 2019). It is plausible that αSyn loss-of-function can induce such a pattern, however further investigation is necessary to elucidate mechanistic insight.

The mechanism of how αSyn regulates DNA DSB repair is still an area for investigation. Our previous work uncovered a novel role for αSyn in the recruitment of 53BP1 to ribosomal DNA DSBs, downstream of γH2AX signaling and upstream of MDC1 activity, in the SK-Mel28 melanoma cell line (Arnold et al., 2024). Furthermore, αSyn has been implicated in regulating DSB repair through a DNA-PK-dependent manner in Hap1 cells (Rose et al., 2024). These data suggest that αSyn may modulate the NHEJ repair pathway, where both 53BP1 and DNA-PK are important. However, the choice between NHEJ and HR is particularly interesting and at the intersection of neurodegeneration and cancer. NHEJ is thought to be the primary DSB repair pathway in post-mitotic cells, like neurons, since it does not require a sister chromatid to act as a template, yet is more error prone (Shrivastav et al., 2008). In contrast, there is growing evidence that different cancers rely primarily on the error-free HR to counteract the genomic instability associated with replicative stress (Helleday, 2010). Studies have shown that a high frequency of melanoma patients harbor mutations in HR-associated genes (Fang et al., 2013, Kim et al., 2021, You et al., 2022), making these tumors vulnerable to immunotherapies and treatments that target HR (Chan et al., 2021, Kim et al., 2021, You et al., 2022). Yet, the choice between HR and NHEJ is a growing topic in the field (Brandsma and Gent, 2012). In our data, αSyn loss-of-function resulted in decreased γH2AX intensity and foci, sex-dependent differences in RPA32 intensity (increased in males, decreased in females), and an increase in nuclear 53BP1 (more robust response in females). This potentially suggests that αSyn upregulation in the TG3 melanoma mouse line is important for functional DDR in a sex dependent way and that when αSyn is no longer present and there is a buildup of unrepaired DNA breaks, male mice can better upregulate HR machinery (RPA32) in the surviving cells, while female mice can better upregulate NHEJ machinery (53BP1) in the surviving cells to try to compensate. Further investigation is necessary to uncover the specific mechanism of how αSyn is influencing the DDR as a function of melanomagenesis and sex *in vivo*.

The sex differences we detect (in tumor onset, growth, and DDR components) are interesting since male sex is a recognized risk factor to the prevalence and outcome of both PD and melanoma. In PD, the prevalence is twice as high in males compared to females and is frequently associated with earlier disease onset (Elbaz et al., 2002a, Moisan et al., 2016); men may develop a postural instability-dominant phenotype, which includes freezing of gait and falling (Elbaz et al., 2002a, Reekes et al., 2020); and men experience more sleep and cognitive issues associated with the disease, such as REM sleep behavior disorder (Reekes et al., 2020) and mild cognitive impairment with a more rapid progression to dementias (Cholerton et al., 2018, Liu et al., 2015). In melanoma, men have a higher risk of developing melanoma across all ages and ethnicities (Scoggins et al., 2006); men exhibit a higher risk of melanoma progression and metastasis than females (Joosse et al., 2011, Scoggins et al., 2006), with a greater risk of mortality (Joosse et al., 2013, Joosse et al., 2011, Smith et al., 2021); and pathologically, thicker and more ulcerated tumors were observed in men (Joosse et al., 2012). In both diseases, there have been many hypotheses as to what is driving these sex differences, including the involvement of sex hormones, the immune system response, and potential environmental exposures. However, in the context of our study, it is interesting to note previously reported sex disparities in DDR pathways. Others have found a greater accumulation of somatic mutations in male cells compared to female cells (Podolskiy et al., 2016), suggesting decreased DNA damage repair. Females have an increased capacity to repair DNA damage by base excision repair (BER) compared to males in mice (Winkelbeiner et al., 2020). Additionally, analysis of molecular difference in 13 cancers from The Cancer Genome Atlas database revealed that DNA repair genes are expressed at higher levels in female patients (Yuan et al., 2016). Furthermore, steroid hormones can regulate DSB repair, both NHEJ and HR (Carusillo and Mussolino, 2020).

Specifically, androgen receptors stimulate the activity and expression of DNA-PK in the NHEJ pathway (Mohiuddin and Kang, 2019), estrogens positively regulate the expression of NBS1 (Komatsu, 2016), and steroid hormones can regulate HR (Bowen et al., 2015, Wilk et al., 2012). Our tumor growth and immunofluorescence data suggest that αSyn plays a role in modulating DSB repair pathways in a sex-dependent manner. Females may be better at upregulating compensatory mechanisms to counteract the unrepaired DNA damage (53BP1 upregulation), and therefore are more resistant to negative effects of αSyn loss-of-function in tumor onset and growth phenotypes. Males may be more vulnerable to DNA damage dysregulation as a consequence of αSyn loss-of function and serve as a more appropriate candidate for therapeutics that target αSyn in melanoma treatment regiments. For example, our data showed a direct relationship between RPA32 increase and *p16* mRNA senescence marker upregulation in male mice and our *in vitro* data, in the male human melanoma cell line, SK-Mel28, αSyn KO significantly impaired growth phenotypes (Arnold et al., 2024).

In summary, the newly generated mouse model, TG3+/+*Snca*-/-, allows for the investigation of the function of αSyn in malignant melanoma. It is possible that individuals with upregulated expression of αSyn may predispose them to Lewy body aggregation in neurons (Osterberg et al., 2015, Schaser et al., 2019), but also melanocytic transformation and melanoma progression. The resulting loss-of-function due to αSyn aggregation (in PD) or gain-of-function of αSyn by increased expression without aggregation (in melanoma) would have differential effects on DNA damage repair pathways, potentially contributing to either neuronal cell death or melanoma cell growth, respectively. Our findings demonstrate the impact of αSyn on melanoma onset, progression, and metastasis in a sex-dependent manner and provide novel therapeutic targets focused on reducing αSyn-mediated DNA repair in melanoma.

## MATERIALS AND METHODS

### Mice

The transgenic TG3 mice(Chen et al., 1996, Pollock et al., 2003, Zhu et al., 2000, Zhu et al., 1998), were established at the Department of Chemical Biology, Rutgers University, Piscataway, USA and provided by Dr. Suzie Chen. Alpha-synuclein KO mice (C57BL/6N- *Sncatm1Mjff*/J) were obtained from Jackson Laboratories (strain #016123). Homozygous alpha-synuclein KO mice were crossed with TG3 heterozygous mice and double heterozygote F1 mice were crossed to each other to generate F2 mice for analysis. Genotyping of mice was carried out by Transnetyx Inc. and primer sequences and protocols are available upon request. For all analyses, homozygous transgenic TG3 *Snca*+/+ and TG3 *Snca*-/- animals (litter mates) were used. Mice were housed in OHSU’s Department of Comparative Medicine (DCM) facilities in a light-dark cycle vivarium. Animals were maintained under *ad libitum* food and water diet. All animal protocols were approved by OHSU IACUC, and all experiments were performed with every effort to reduce animal lives and animal suffering, according to the NIH Guide for the Care and Use of Laboratory Animals.

### Tumor growth analysis

Starting at P30, mice underwent isoflurane anesthesia every 10 days to assess weight and tumor growth. Researchers were blinded to condition. To quantitate the severity of melanoma progression, detailed observation and photodocumentation was used to assign numerical scores of 0 to 5 to the thickness of pinna tumors (see Figure S2 for detailed description) or quantitative measurements for thickness of perianal tumors. For pinna tumors, 0=tumor not palpable or visible; 1=individual small, clearly recognizable nodes or elevations in skin; 2=small, numerous recognizable nodes or elevations; 3=significantly thickened ears, clearly nodular tumors; 4=severely thickened ears or coarse tumors; 5=extreme tumor growth with risk of ulceration. Tumor onset was designated as time when tumor changed from 0 to 1. For perianal tumors, a ruler was used to manually measure the length of the perianal tumor in centimeters.

### Immunofluorescence staining

For immunofluorescence staining of perianal tumors, 5μm sections of formalin-fixed and paraffin-embedded tissue blocks were deparaffinized and bleached in a H2O2 solution for 1 hour at room temperature (1% dipotassium phosphate, 0.5% potassium hydroxide, 3% hydrogen peroxide). Tissue underwent antigen retrieval overnight at 56C (10 mM Tris base, 1mM EDTA solution, 0.05% Tween 20, pH 9.0). Samples were permeabilized in 0.25% Triton X-100 in PBS for 10 minutes and blocked in 2% FBS/1% BSA in PBS for 2 hours and then placed in the primary antibody overnight at 4C. The next morning, samples were washed in 1x PBS and placed in secondary antibody for 1 hour at 37C. Samples were washed 4 times in 1x PBS. The third wash contained DAPI (2.5µg/ml) for 20min. Coverslips were mounted using CFM2 antifade reagent and sealed with BioGrip. All immunofluorescence images were taken on a Zeiss Laser-Scanning Confocal Microscope 980 with Airyscan and analyzed in Arivis Software. Mean intensity was measured after imposing DAPI masks over each nucleus. All cells within a 63x image were analyzed and numbers of n are provided in each figure legend. Statistical significance was assigned using T-test.

Antibody specifics were as follows: LB509 (Abcam #27766, RRID:AB_727020, 1:500), RPA32 (Bethyl #A300-245A, RRID:AB_210547, 1:1000), γH2AX (Cell Signaling #9718, RRID:AB_2118009, 1:500), 53BP1 (BD Biosciences #612522, RRID:AB_2206766, 1:1000).

### Quantitative RT-PCR

Pinna tumors, perianal tumors, and lymph nodes were homogenized in RNeasy mini kit buffer (Qiagen) using a hand-held tissue homogenizer followed by Qiashredder centrifugation (Qiagen). Total RNA was isolated using the RNeasy mini kit (Qiagen) according to the manufacturer’s instructions. RNA concentration was measured with a NanoDrop spectrophotometer and cDNA was synthesized from 500ng RNA with M-MLV reverse transcriptase (Promega). Analysis of mRNA expression was performed using quantitative Real-Time PCR on the QuantStudio 3 (Applied Biosystems). A volume of 1 μl cDNA template, 1 μl of forward and reverse primers (each 10 μM) and 10 μl of SYBR Green I (Roche) were combined to a total volume of 20 μl. Primers used are described in Table S1. Each sample was analyzed in duplicates. The target cDNA was normalized to β-Actin levels. Statistical significance was assigned using T-test.

### Statistical analysis

Beyond individualized analysis within each assay methodology, all data was processed using GraphPad Prism version 9.0. Data was analyzed using T-test, unless stated otherwise, and considered statistically significant if p < 0.05. All data was presented as a mean +/- standard error of the mean (SEM).

## Abbreviations

Alpha-synuclein: αSyn
Parkinson’s Disease: PD
DNA damage response: DDR
knockout: KO
double-strand break: DSB

## DATA AND MATERIALS AVAILABILITY

All data are available in the main text or the supplementary materials. No datasets were generated or analyzed during the current study.

## ACKNOWLEDGEMENTS

All immunofluorescence imaging could not have been possible without help from Stefanie Kaech Petrie, Felice Kelly, and Hannah Bronstein in the OHSU Advanced Light Microscopy Core (RRID:SCR_009961). We thank Dr. Pamela Cassidy (OHSU) for the invaluable discussions. Thank you to Dr. Suzie Chen (Rutgers University) for providing the TG3 mouse line and help with genotyping protocols. We thank Dr. Sebastian Staebler and Dr. Anja Bosserhoff (Friedrich-Alexander University of Erlangen-Nurnberg) for the invaluable advice and help with TG3 mouse line management, qRT-PCR protocols, and IF protocols. Lastly, many thanks to Unni Lab members who helped with the revision process: Dr. Carlos Soto Faguas, Dr. Elizabeth Rose, Anna Bowman, Elias Wisdom, Jessica Keating, and Samuel Kim.

## Funding

The work was supported by:

National Institute on Aging of the National Institutes of Health F30AG082406

National Institute of Health R01NS102227

National Institute of Health P30NS061800

National Institute of Health P30CA065823

Melanoma Research Alliance

Michael J Fox Foundation

Kuni Foundation

Melanoma Research Foundation Medical Student Research Award

Oregon Partnership for Alzheimer’s Research

## AUTHOR CONTRIBUTIONS

Conceptualization: MRA, VKU

Data Curation: MRA, VKU

Formal Analysis: MRA, VKU

Funding Acquisition: MRA, VKU

Investigation: MRA Methodology: MRA, VKU

Project Administration: MRA, VKU

Resources: MRA, SC, VKU

Software: MRA, VKU

Supervision: MRA, VKU

Validation: MRA, VKU

Visualization: MRA

Writing- original draft: MRA, VKU

Writing- review & editing: MRA, SC, VKU

---

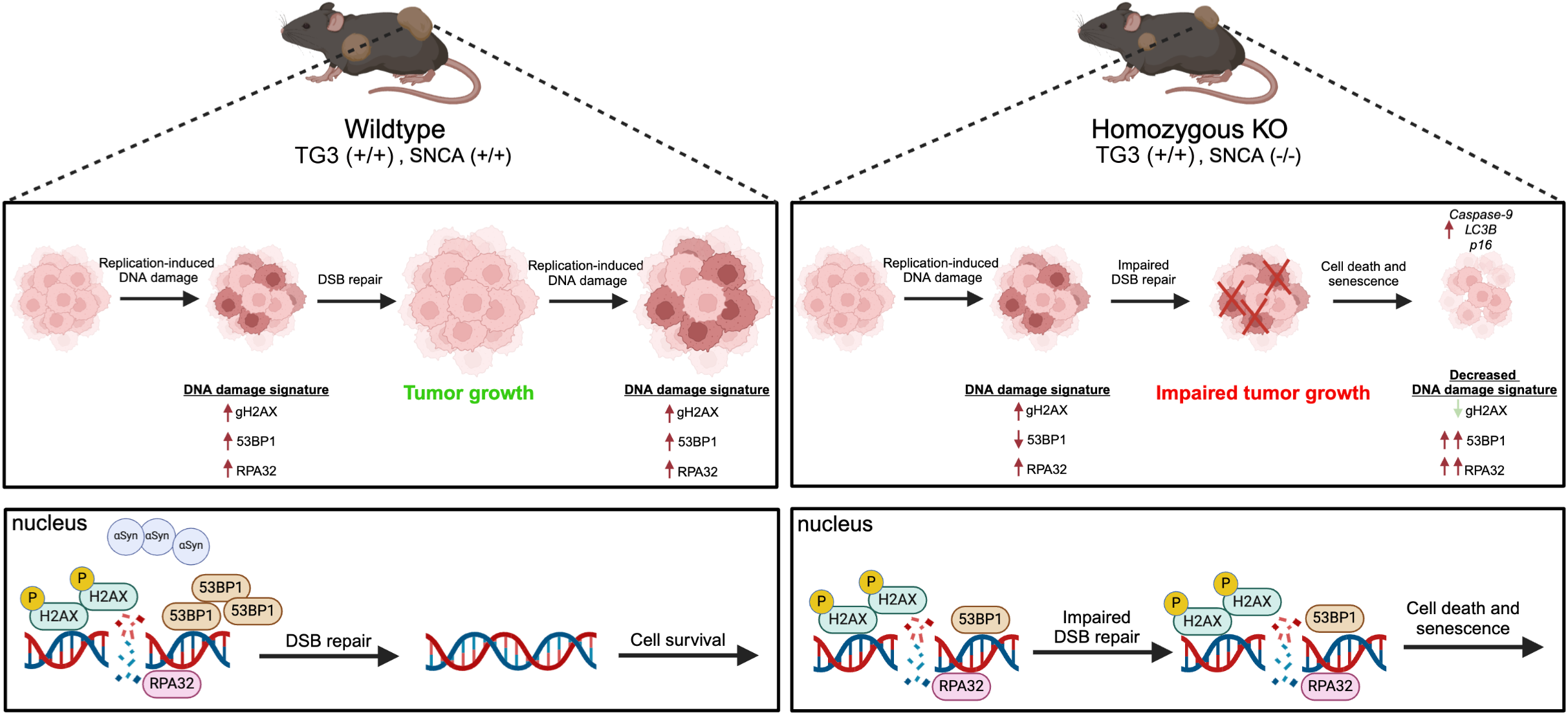

