## Supplemental Materials for "Alpha-synuclein knockout impairs melanoma development and alters DNA damage repair in the TG3 mouse model in a sex-dependent manner"

Moriah R. Arnold *et al.*

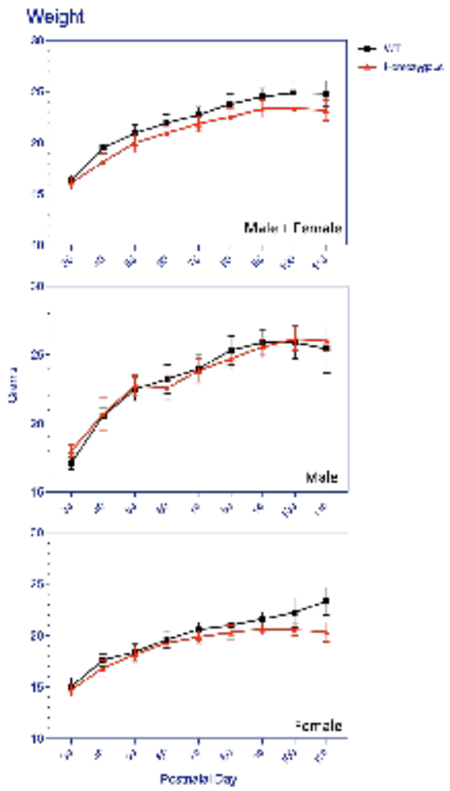


**Figure S1. Alpha-synuclein knockout does not affect mouse weight.**
Weight of TG3+/+*Snca*+/+ (n=15) and TG3+/+*Snca*-/- (n=14) from P30 to P110 (endpoint). Analysis was further stratified by sex with TG3+/+*Snca*+/+ male (n=10), TG3+/+*Snca*+/+ female (n=5), TG3+/+*Snca*-/- male (n=7), and TG3+/+*Snca*-/- female (n=7). Error bars represent Standard Error of the Mean (SEM). Statistical testing by two-way ANOVA. Weight is represented in grams.

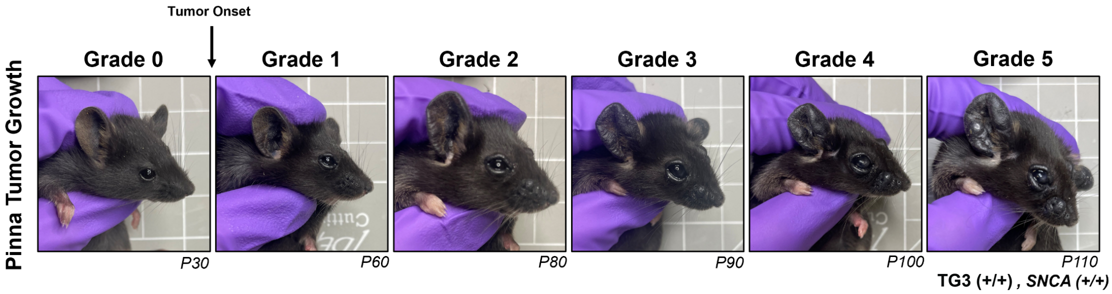


| Grade | Description |
|---|---|
| 0 | Tumor not palpable or visible |
| 1 | Individual small, clearly recognizable nodes or elevations in skin |
| 2 | Small, numerous recognizable nodes or elevations |
| 3 | Significantly thickened ears, clearly nodular tumors |
| 4 | Severely thickened ears or coarse tumors |
| 5 | Extreme tumor growth with risk of ulceration |

**Figure S2. Representative images and description of pinna tumor grading scale.**

| Primer | Forward (5'-3') | Reverse (5'-3') |
|---|---|---|
| β-Actin | TGGAATCCTGTGGCATCCATGAAAC | TAAAACGCAGCTCAGTAACAGTCCG |
| Grm1 | GGGCAGGGAACGCCAATTCT | TGGAAGGGCTGCTGGGAGGG |
| Caspase-3 | AGCAGCTTTGTGTGTGTGATTCTAA | AGTTTCGGCTTTCCAGTCAGAC |
| Caspase-9 | TCCTGGTACATCGAGACCTTG | AAGTCCCTTTCGCAGAAACAG |
| RIP3 | AAGTGCAGATTGGGAACTACAACTC | AGAATGTTGTGAGCTTCAGGAAGTG |
| LC3B | CCCCACCAAGATCCCAGT | CGCTCATGTTCACGTGGT |
| Cdkn2a (p16) | CCCAACGCCCCGAACT | GCAGAAGAGCTGCTACGTGAA |

**Table S1. Primers used in qRT-PCRs.**
